# A global effort to benchmark predictive models and reveal mechanistic diversity in long-term stroke outcomes

**DOI:** 10.1101/2024.10.17.618691

**Authors:** Anna Matsulevits, Pedro Alvez, Manfredo Atzori, Ahmad Beyh, Maurizio Corbetta, Federico Del Pup, Lilit Dulyan, Chris Foulon, Thomas Hope, Stefano Ioannucci, Gael Jobard, Hervé Lemaitre, Douglas Neville, Victor Nozais, Christopher Rorden, Orionas-Vasilis Saprikis, Igor Sibon, Christoph Sperber, Alex Teghipco, Bertrand Thirion, Louis Fabrice Tshimanga, Roza Umarova, Ema Birute Vaidelyte, Emiel van den Hoven, Esteban Villar Rodriguez, Andrea Zanola, Thomas Tourdias, Michel Thiebaut de Schotten

## Abstract

Stroke remains a leading cause of mortality and long-term disability worldwide, with variable recovery trajectories posing substantial challenges in anticipating post-event care and rehabilitation planning. To address these challenges, we established the NeuralCup consortium to benchmark predictive models of stroke outcome through a collaborative, data-driven approach. This study presents findings from 15 international teams who used a comprehensive dataset including clinical and imaging data, to identify and compare predictors of motor, cognitive, and emotional outcomes one year post-stroke. Our analyses integrated traditional statistical approaches and novel machine learning algorithms to uncover ’optimal recipes’ for predicting each domain. The differences in these ‘optimal recipes’ reflect distinct brain mechanisms in response to different tasks. Key predictors across all domains included infarct characteristics, T1-weighted MRI sequences, and demographic factors. Additionally, integrating FLAIR imaging and white matter tract analysis significantly improved the prediction of cognitive and motor outcomes, respectively. These findings support a multifaceted approach to stroke outcome prediction, underscoring the potential of collaborative data science to develop personalized care strategies that enhance recovery and quality of life for stroke survivors. To encourage further model development and validation, we provide access to the training dataset at http://neuralcup.bcblab.com

## 1. Introduction

Knowledge about a future event empowers individuals to adequately prepare and take measures to influence outcomes in their favor. In healthcare, mathematical and statistical modeling has enabled the use of past observations to anticipate outcomes following various life-changing events such as brain damage. Stroke, as the second leading cause of death, has seen a 70% increase in incidence in the past two decades^1^, resulting in 6.55 million deaths and 101 million survivors in 2019^1^. While recanalization of the occluded vessel mitigates damage of brain tissue and is therefore associated with a relief of post-stroke symptoms, less than 10% of acute stroke patients actually receive this highly effective treatment (e.g. thrombectomy) due to strict eligibility criteria^2^. Despite recent tremendous progress with recanalisation interventions, many of the affected individuals will continue to have persistent motor and cognitive deficits that impact their quality of life, their ability to communicate and socialize, as well as their ability to return to work^3^. Hence, predictive frameworks in stroke are invaluable for providing realistic prognostic expectations to the patients and their families, accurately planning post-acute care and enhancing clinical research trials by identifying patients with an expected homogenous outcome. This will allow to counter-balance individuals with similar outcome trajectories in clinical trials, increasing the statistical power of such studies^4^.

Forecasting behavioral outcomes post-stroke is a longstanding challenge^5^, but the optimal predictive biomarkers, neuroimaging methodology, or algorithms are difficult to pinpoint due to significant variations in study resources, methods, and purpose. Neuroimaging has been promising in revealing brain alterations due to infarct characteristics, particularly helping individual prediction^6,7^. Previous work has demonstrated the importance of neuroimaging metrics such as stroke volumes^8^, stroke location^9^, disconnection^7^, functional pattern^10^, co-existence of small vessel disease^11^, and pre-existing brain atrophy^12^. These insights influence prediction parameters (for a review, see^14,15^), though methodologies vary from conventional regression models^16^ to advanced machine learning algorithms^7^. However, few frameworks consider cognitive and neuropsychological outcomes on top of the more visible motor outcomes^17^, and most focus on 3-month outcomes, limiting long-term outcome investigations. In addition, many studies mislabel statistical associations as ‘predictions’^18,19^ which do not forecast accurate outcomes in new data. Advancing precision medicine requires data-driven approaches that assess model performance on unseen data to predict outcomes in new cohorts^20^.

Recently, systematic reviews have compared outcome prediction^21,22^ but have often excluded machine learning algorithms and failed to provide comprehensive comparisons of available methodological approaches and their combinations. Yet, limited efforts place their existing predictive methods into a comparative context using the same dataset.

To address these gaps, we organized an open competition involving 15 teams of experts in stroke outcome prediction from across the globe at the NEURAL conference. Participants used a comprehensive dataset, including motor, cognitive, and neuropsychological scores one year after stroke, as well as neuroimaging, lesion, and demographic data to predict outcomes on an unseen out-of-sample dataset. Following this, we conducted a statistical analysis of the input and method combinations to derive the best recipe for predicting a wide range of stroke outcomes. To encourage further model development and validation, we provide access to the training dataset at http://neuralcup.bcblab.com.

## 2. Methods

### Dataset

The dataset used for predicting stroke outcomes on three different domains – motor, cognitive, psychological – is based on a prospective observational study, “Brain Before Stroke (BBS)” cohort, conducted between June 2012 and February 2015. The main results and secondary objectives have already been published^16,23–25^. The study was approved by the local research ethics committee, and written consent was obtained from the patients before inclusion. The BBS cohort has not been previously used to compare different prognosis models side-by-side, which is the objective of the present study.

Briefly, the BBS cohort recruited consecutive stroke patients who underwent clinical and MRI evaluations at initial (24 to 72 hours) and chronic (1 year) states. Primary inclusion criteria were men and women over 18 years old with a clinical diagnosis of minor-to-severe supratentorial cerebral infarct (NIHSS between 1 and 25) at the stroke onset. Exclusion criteria were history of symptomatic cerebral infarct with a functional deficit (pre-stroke modified Rankin Scale [mRS] score ≥1), infratentorial stroke, history of severe cognitive impairment (dementia), or psychiatric disorders according to axis 1 of the DSM-IV^26^ criteria except for major depression, coma, pregnant or breast- feeding women, and contraindications to MRI.

We mainly aimed to compute and compare imaging modalities’ predicting abilities but also included demographic data such as age and sex, knowing their predictive value^27^. The baseline imaging used for prediction consisted of MRI performed between 24 and 72 hours after stroke onset on a 3T scanner (Discovery MR750w, GE Healthcare). We provided the raw Diffusion-Weighted Images (DWI) and the infarct masks that had already been manually delineated^25^ regarding the critical role of stroke volume and location in the outcome^17,28^. Additionally, we included 3D FLAIR and 3D T1 sequences, which, while capable of detecting stroke-related changes at this stage, offer valuable insights into the overall integrity of the brain beyond the lesion site^29^. Furthermore, the teams were free to use any openly available brain atlases^30,31^ or additional tools to compute supplementary inputs such as the white matter tracts and/or their disconnection^32–35^ by the infarct. In summary, each team was free to include any input combination in their predictive models, which resulted in the following 9 different inputs: “age”, “gender”, “DWI”, “T1”, “segmented lesions”, “FLAIR”, “tracts”, “parcellation atlases”, and “disconnectome”. The parameters of acquisition and additionally used inputs are summarized in Table 1.

**Table 1.**
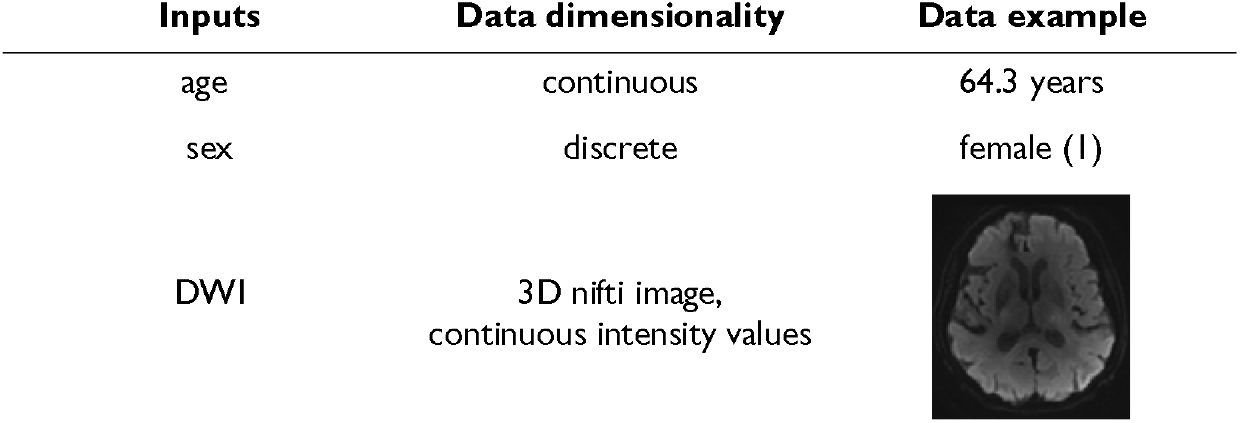
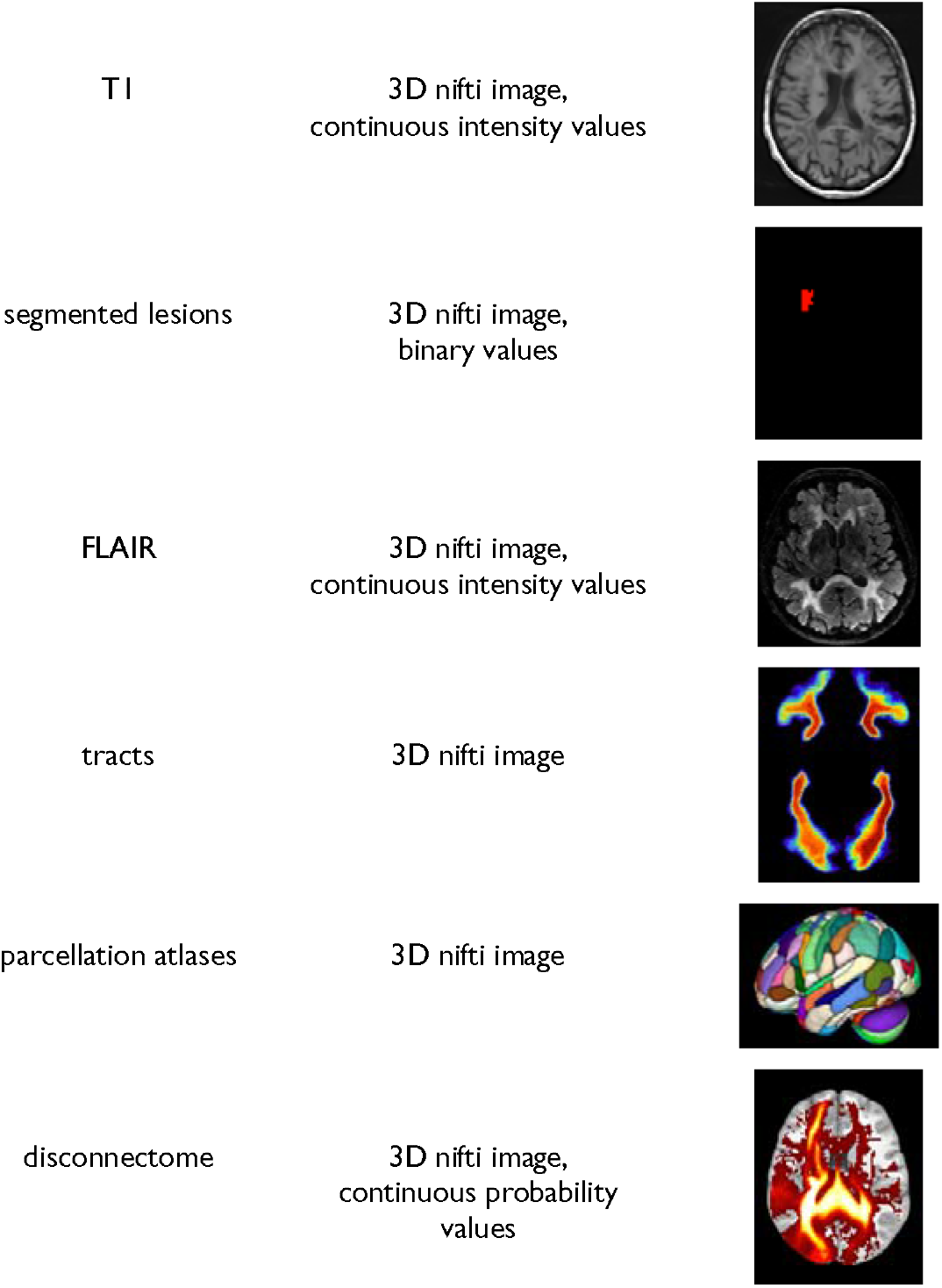
Inputs provided for the outcome prediction, as well as additional inputs (tracts, parcellation atlases, disconnectome) used by the teams.

| Inputs | Data dimensionality | Data example |
| --- | --- | --- |
| age | continuous | 64.3 years |
| sex | discrete | female (1) |
| DWI | 3D nifti image,<br>continuous intensity values |  |
| T1 | 3D nifti image,<br>continuous intensity values |  |
| segmented lesions | 3D nifti image,<br>binary values |  |
| FLAIR | 3D nifti image,<br>continuous intensity values |  |
| tracts | 3D nifti image |  |
| parcellation atlases | 3D nifti image |  |
| disconnectome | 3D nifti image,<br>continuous probability<br>values |  |

For 1-year post-stroke, teams had to predict motor, cognitive, and psychological outcomes. We intentionally selected the 1-year data-subset considering this time point reflects a more permanent level of handicap rather than the fluctuating recovery seen in earlier months. Motor outcome was based on the full Fugl-Meyer (FM) score (range: 0–242), covering five domains: motor function, sensory function, balance, passive joint range of motion, and joint pain during passive movement. The score provides a detailed evaluation of upper and lower limb functions with an excellent inter and intra-rater reliability^36^. We used the total score (FM total), as well as specifically the motor domain (FM motor). Cognitive outcome was assessed with the Montreal Cognitive Assessment (MoCA) which is a validated tool for screening cognitive impairment after stroke^37^ (range: 0–30). To assess more details on the affected domains, we also considered the sub-scores of the MoCA (Attention, Visuospatial, Denomination, Language, Abstraction, Orientation, Recall) and the Isaacs test set (IST) which evaluates the executive functions through categorical verbal fluency^38^ (range: 0–max. number of orally produced words for different categories within 1 minute). Psychological outcome was assessed using the Hospital Anxiety and Depression Scale (HAD-Anxiety and HAD-Depression)^39^ (range: 0–21 for each test).

From the original BBS dataset, we excluded individuals with inadequate baseline neuroimaging (missing sequences, insufficient image quality, no stroke at MRI, infratentorial ischemic stroke) or missing 1-year values on the scores to predict, resulting in 237 patients (see Supplementary Figure 1 for a detailed flowchart). The remaining data was split into a training dataset (n=187) and a validation dataset (n=50) through pseudorandomization to ensure comparable distribution of the outcome scores (Supplementary Figure 2). Given the time (one year post-stroke) of assessment, most patients have fairly but not fully recovered from their motor and cognitive impairment, leading to scores indicative of rather mild symptoms. However, these impairments remain handicapping for the individuals’ everyday life as more than 70% of patients employed before stroke do not return to work after the incident ^40,41^. Table 2 provides patients’ demographics and neuropsychological scores of the whole dataset.

**Table 2.** Patient demographics and average scores (with standard deviations) for the training and validation cohort.

| Score | training |  | validation |  |
| --- | --- | --- | --- | --- |
| | Mean (x±) | SD | Mean (x±) | SD |
| Age | 65,44 | 13,48 | 66,29 | 12,61 |
| FM | 224,93 | 30,43 | 224,66 | 31,16 |
| MoCA | 24,13 | 4,93 | 24,38 | 5,78 |
| IST | 30,48 | 6,41 | 30,66 | 9,23 |
| HAD-A | 6,07 | 3,80 | 5,78 | 4,27 |
| HAD-D | 4,54 | 3,98 | 4,10 | 3,03 |

**Note:** FM (Fugl-Meyer), an expanded version of the score was used covering the range (0-242), which includes five evaluation domains (motor function, sensory function, balance, passive joint range of motion, and joint pain during passive movement), lower score indicating more motor impairment; MoCA (Montreal Cognitive Assessment) with a range of 0-30, lower score indicating more cognitive impairment; IST (Isaacs test set) ranging from 0 to the number of words produced, lower score indicating less fluency and impaired executive functions; HAD-A (the Hospital Anxiety and Depression Scale – Anxiety); HAD-D (the Hospital Anxiety and Depression Scale – Depression), both with a range of 0-21, higher score indicating more anxiety and depression.

The participating teams were provided with data and instructed to produce clinical score predictions for each of the 50 out-of-sample patients. There were no restrictions placed on the use of inputs or methods. The teams’ performances were primarily ranked based on the overall Pearson mean R^2^ outcome^42^, but we also examined additional statistical outcomes such as the Mean Squared Error (MSE) and Mean Absolute Error (MAE) losses (for formulas see Supplementary Formula 1 and Formula 2). To assess clinical relevance, we also used the area under the receiver operating characteristic (ROC) curve for motor, cognitive, and psychological tests (FM, MoCA, IST, HAD-A, and HAD-D). With clinically validated thresholds to define poor outcomes (FM ≤ 100, MoCA ≤ 25, IST ≤ 28, HAD-A ≥ 8, HAD-D= ≥ 8), we categorized the individual patient predictions of the teams and evaluated these against the actual category the individuals fell into.

### Exhaustive evaluation of the used approaches

After evaluating teams’ performances, we aimed to quantify which combinations of inputs and methods led to the most accurate stroke outcome prediction. The teams utilized different methods which were categorized into eight main classes that represent different strategies to obtain the prediction *(Artificial Neural Networks, Regression)*, to extract and represent the data (*Clustering, Feature Selection, Dimension Reduction, Parcellation/Segmentation),* and to validate the prediction *(Cross Validation, Bootstrapping)*. More details on each class, alongside examples of specific techniques used by the participating teams, can be found in Supplementary Table 1.

To explore the similarities and differences between the teams we used Uniform Manifold Approximation and Projection for Dimension Reduction (UMAP)^43^ on a matrix summarizing the observed feature combinations and obtained R^2^ for each score. This created a 2D discrete morphospace – 60x60 grid to allow statistics on the UMAP latent space – where teams were localized based on the combinations of inputs (among the nine listed in Table 1) and methods (among the eight listed in Supplementary Table 1) they used. Similar approaches were located close to each other while differing methods were further apart. The low dimensionality of the space allowed us to utilize the matrix for the next analysis step. Following previous work^5^, using the FSL tool *randomise*^44^ we identified areas in the morphospace that were associated with a high R^2^ score for each test. The *randomise* analysis yielded t-statistic maps distinguishing significant from non-significant regions, marking the presumably ‘best’ prediction points that we defined based on the local maxima. With the UMAP inverse transformation (*inverse transforms*) it is possible to generate a high dimensional data sample given a location in the low dimensional embedding space, meaning we can obtain an input data vector from coordinates in the morphospace even for coordinates that are not corresponding to the initial input data points. Therefore, we then used the UMAP inverse for the combination of variables that were assigned to the points where the t-value for the *randomise* test was the highest, associated with the best-predicting performance. After applying the inverse option of the analysis, we obtained a separate matrix for each feature, each containing 3600 values (60x60 space). Every value in the matrix represented a point in the UMAP space, corresponding to one out of seventeen (total number of features: 8 inputs + 9 methods) components of a potential feature vector. We binarized each feature matrix using the threshold 0.5 and concatenated the seventeen feature matrices that summarize the inputs and methods we were investigating. Subsequently, we were able to obtain a combination of used and discarded features for the previously identified local maxima (as well as any other coordinate in the space), revealing an estimation of the theoretically best combination for accurately predicting the one-year outcome for each clinical domain (see Figure 1).

**Figure 1:**
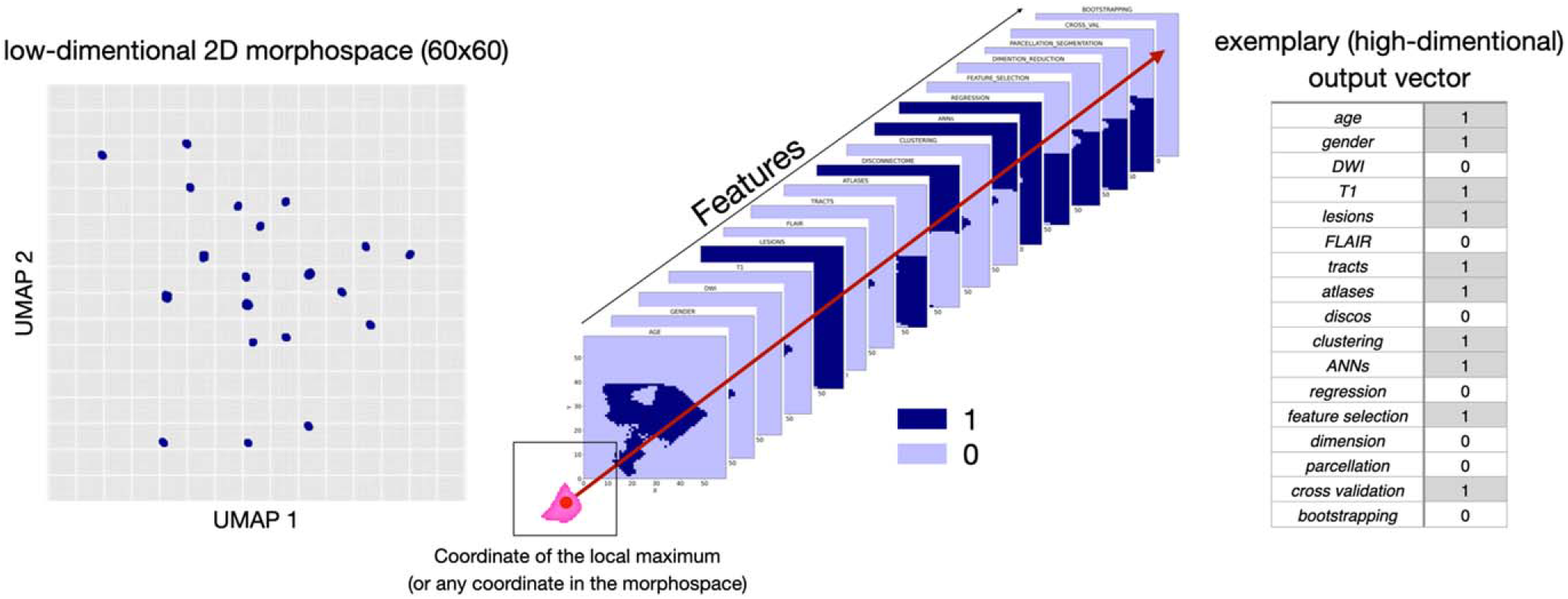
Visualization of the process for obtaining the theoretically optimal feature combination for predictions: inversing the randomise analysis yields heatmaps for each analyzed feature. After binarizing these maps, we investigated the overlap of the local maxima of the clinical tests with the binarized feature maps (1 representing the presence of the feature, and 0 representing the absence of a feature in the final combination ‘optimal recipe’).

## 3. Results

### Patient characteristics, participating teams and their prediction approaches

Fifteen teams from nine different countries, each with different scientific backgrounds, participated in the stroke outcome prediction and submitted 24 stroke outcome predictions in total (Figure 2a). They combined between none and eight input data with one to five methods.

**Figure 2:**
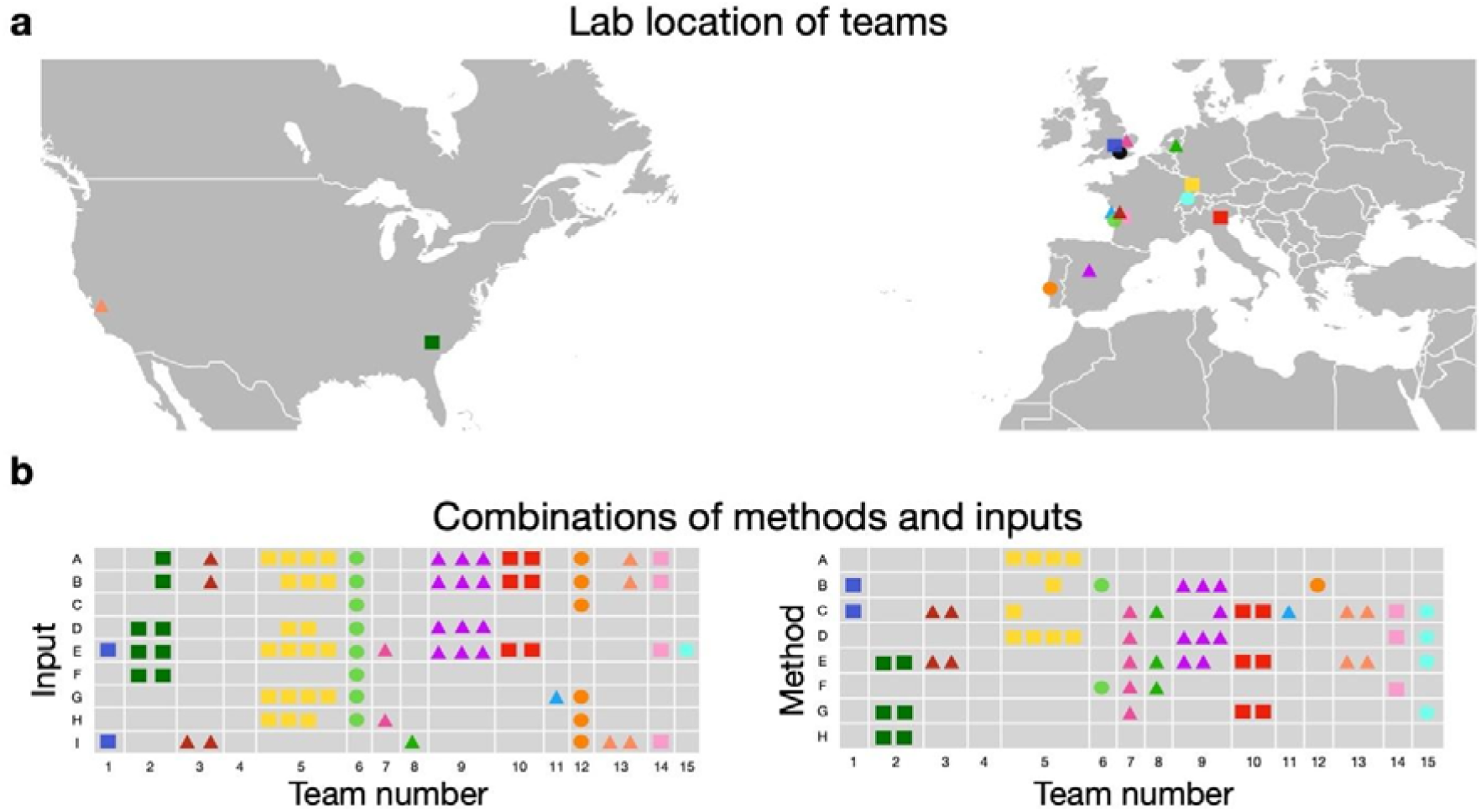
Summary of participating teams and the approaches taken for all predictions. **a** Locations of the teams’ affiliated labs. **b** Summary of different inputs (A: age, B: gender, C: DWI, D: T1, E: lesion, F: FLAIR, G: tracts, H: atlases, I: disconnectome) and methods (A: clustering, B: artificial neural networks, C: regression, D: feature selection, E: dimensionality reduction, F: parcellation, G: cross-validation, H: bootstrap) used for each prediction. (Figure modified from^38^).

### Comparison of the evaluation metric R^2^

The performance of each team’s model was evaluated by comparing the predicted scores for the 50 out-of-sample patients with their actual recorded scores. The mean proportion of variance explained (R^2^) was calculated for each of the 24 prediction models (Figure 3a) from the 6 main scores: FM motor, FM total, MoCa, IST, HAD-A, and HAD-D. Team 5, in their third prediction obtained the highest overall R^2^= 0.311, followed by the same team’s first prediction with R^2^ =0.238, and the second prediction of team 2 (R^2^ =0.182). The mean R² values displayed a strong disparity of the predictive performances according to the type of outcome. The prediction of the motor outcome at one year could reach R² as high as 0.611 for team 5. On the other hand, the prediction of the neuropsychological outcome was the worst with R² not higher than 0.034 to anticipate post-stroke depression (HAD-D) and 0.138 for post-stroke anxiety (HAD-A). The predictive performances of cognitive status were intermediate between motor and mood prediction. Exhaustive information describing the results of all teams can be found in the supplementary material (Supplementary Table 2 and Supplementary Table 3).

**Figure 3:**
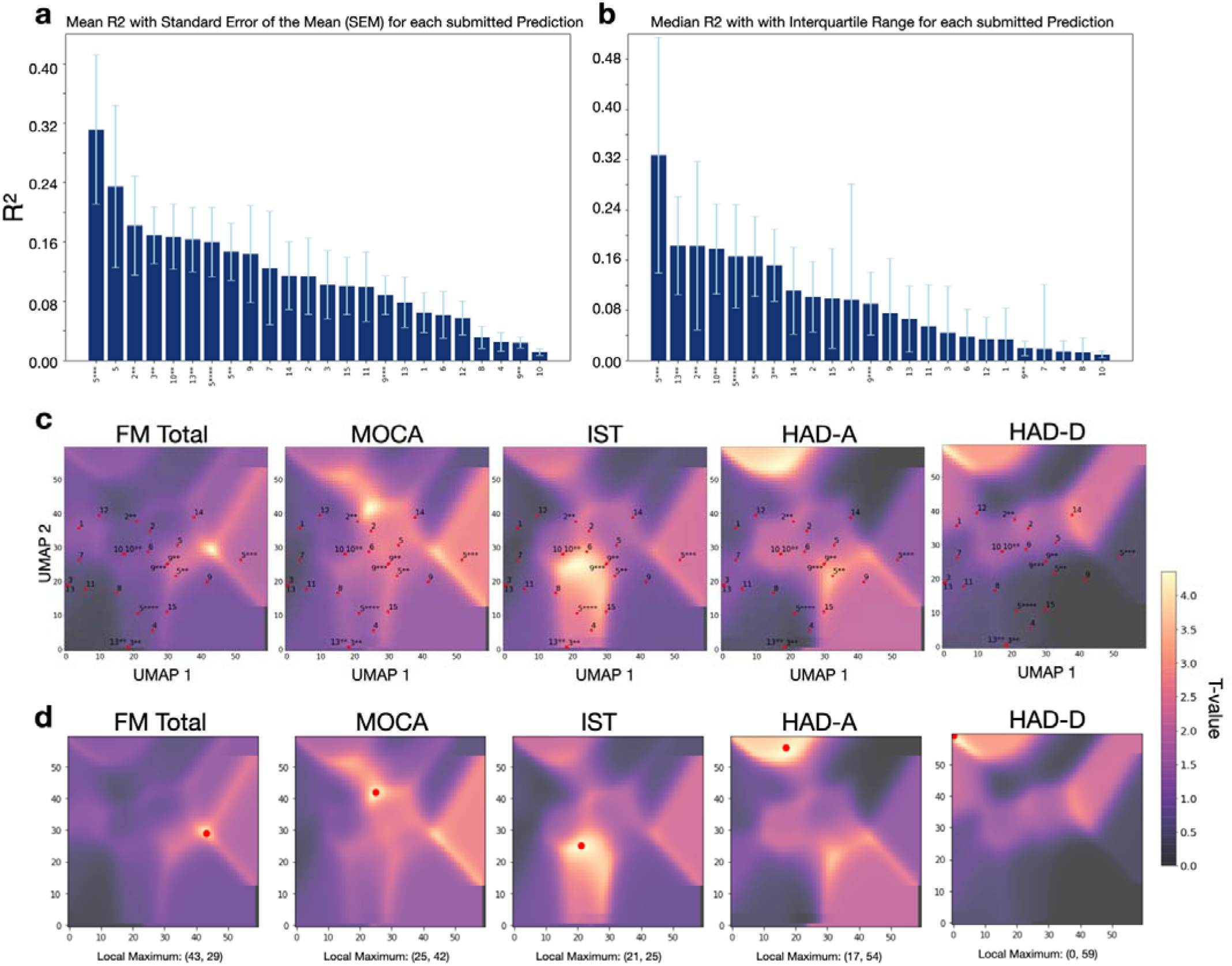
**a** Mean R^2^ comparison for all submitted predictions (motor, cognitive, and psychological outcomes) of five neuropsychological scores (FM total, MoCA, IST, HAD-A, HAD-D) sorted ascendingly from the highest score to the lowest score across all teams whose number is indicated on the x-axis. The stars indicate the prediction number, the whiskers indicate the standard error of the mean (SEM). **b** Median R^2^ comparison for all submitted predictions of the same scores, sorted ascendingly with whiskers indicating the interquartile range (IRQ). **c** T-statistic maps for each clinical outcome test, obtained from the FSL randomise analysis. The yellow regions indicate a significant t-value, the purple regions indicate a non-significant t-value. The UMAP distribution of all teams is plotted on the t-statistic maps. **d** Local maximum of the t-statistic map for each analyzed outcome score.

As secondary analyses, we also looked at how each team’s model could classify patients’ outcomes as poor or good based on clinically relevant thresholds. The ROCs of the five best models are shown in Supplementary Figure 3 and confirmed the highest performances for motor prediction followed by cognitive and mood predictions. As an additional metric for evaluation, we calculated the mean average errors (MAEs) and mean squared errors (MSEs) of the teams’ predictions for the main test scores, which can be found in Supplementary Table 6.

### Evaluation with UMAP and randomise approach

To investigate the nuances of different combinations of features (input variables and methodological approaches) used by the teams, we reduced their interactions into a UMAP morphospace for each outcome. These representations, combined with t-statistic maps (that indicate locations of the space associated with higher R²) highlight how close or far each of the 24 models is from the best prediction. The observed pattern indicated that overall, the proximity of the models’ placement in the morphospace was closer to the local maxima for motor outcomes and cognitive outcomes compared to psychological outcomes (Figure 3c and d). In other words, the prediction models show generally closer distances to the local maxima (indicated through the highest t-value) of the test scores FM, MoCA, and IST compared to the scores HAD-A and HAD-D, which is reflected in the achieved prediction accuracies.

Moving back from the local maximum in the morphospace to the combination of features associated with this location, we identified the optimal combinations of features (defined as the best approaches within the boundaries of the given context) for each of the motor, cognitive, and neuropsychological domains (see Table 3 and Figure 1). The overview reveals that the best predictions always required the lesion mask as an input. Adding information about the overall status of the brain by including the FLAIR modality improved predictions on the cognitive domain whereas utilizing white matter tracts and atlases seem to be beneficial for anticipating motor outcomes. Conversely, predicting higher cognitive outcomes in the neuropsychological domain benefited more from a global disconnectivity analysis.

**Table 3.**
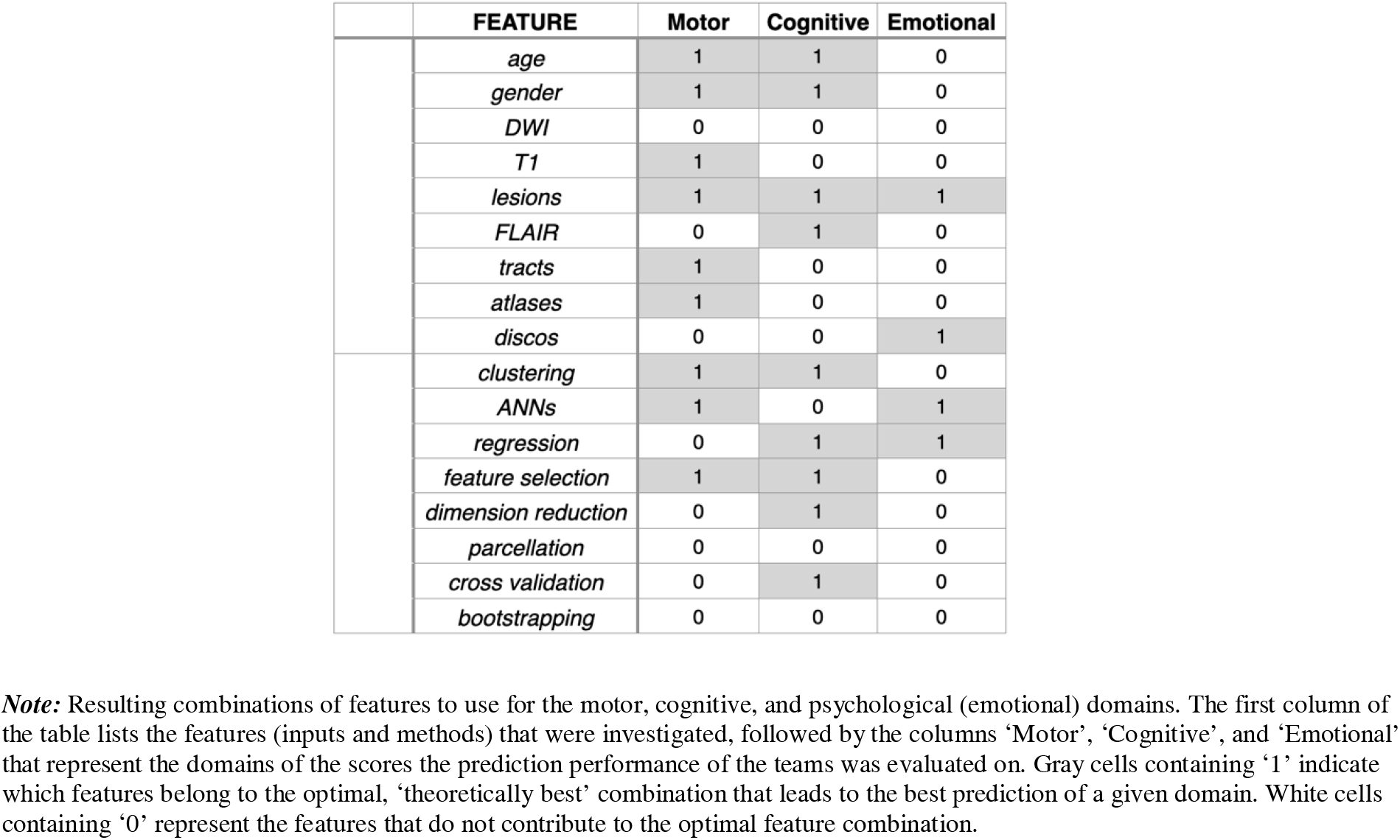
Summary of the optimal combination of features to use for long-term (1 year) predictions of motor, cognitive, and psychological stroke outcomes.

Investigating the methodological approaches, the most prominent and successful methods were clustering, ANNs (Artificial Neural Networks), regression, and feature selection. However, the utilization of the different methods varies (e.g., different clustering), making comparison less straightforward than with inputs. For an exhaustive listing of combinations for each single score of FM, MOCA, IST, HAD-A, and HAD-D, please refer to Supplementary Table 4.

Examining the obtained t-statistics maps from the first step, we can identify separate clusters of significant t-values (Supplementary Figure 4). Repeating the above-described procedure for the local maxima of each cluster separately led to slightly different feature combinations for different clusters of the same test. For instance, the analysis of the local maximum of one cluster (coordinates: 25, 42) that was highly predictive for MoCA resulted in the combination of FLAIR, clustering, feature selection, and dimension reduction while the analysis of the second cluster (local maximum coordinates: 44, 28) resulted in a different combination of features, that included age, sex, T1, lesions, ANNs, regression, and feature selection (Supplementary Figure 4). This example, along with similarly varying results for different significant clusters of the HAD-A and HAD-D tests, demonstrates that there is potentially more than one recipe that leads to a decent prediction of a score. The differences found in the feature combinations for the tests also show that there is not one single recipe for an overall good prediction that covers all stroke outcomes. Instead, different combinations of features work differently well depending on the type of outcome being predicted.

## 4. Discussion

Our study underscores the potential for long-term stroke outcome predictions across multiple domains. By systematically analyzing different feature combinations, we demonstrate that there is no universal solution for predicting stroke outcomes, reinforcing the notion that distinct recovery mechanisms require tailored approaches^45,46^. The effectiveness of predicting models varies significantly across motor, cognitive, or emotional outcomes, necessitating domain-specific methodologies. A key innovation of this study is the use of Uniform Manifold Approximation and Projection (UMAP) with inverse transformation to benchmark predictive models. This framework allows for a novel exploration of interactions between input features and methodologies, providing valuable insights into prediction strategies.

The principal conclusion of this manuscript derives from an in-depth analysis of the methodologies employed by the participating teams in constructing their predictive models and the identification of the features instrumental in facilitating optimal long-term predictions. This consortium has established, to date, one of the most comprehensive out-of-sample stroke outcome prediction comparisons, elucidating distinct ‘optimal recipes’ for various outcome scores, which reflect the different brain mechanisms in response to different tasks. These findings underscore that not all inputs and methodologies exhibit equivalent efficacy in the precise prognostication of specific stroke outcomes. Further, the result highlights the importance of individualized predictions, as tailoring models to specific patient characteristics and incorporating neurovariability is crucial to enhancing predictive accuracy and clinical relevance. With the exhaustive information captured in different neuroimaging data^47^, we are able to unravel an important source of variance, improving our understanding of each patient’s unique brain architecture, and applying personalized strategies to ensure the best care. Lesion-symptom mapping, with its rich history in studying the brain structure-function relationships, remains a widely used approach for analyzing and explaining stroke outcomes^48^. In our case, for predictions of all domains (motor, cognitive, emotional), the location and size of the lesion were consistently identified as pivotal factors.

Some other factors were specific to a domain. For instance, the prediction of cognitive scores, such as the MoCA, was found to derive greater benefit from FLAIR imaging. This modality not only delineates the infarct but also captures underrepresented characteristics of cerebral damage beyond the lesion site, particularly white matter hyperintensities indicative of small vessel disease^49^. Such proxies of brain health^50^, if applied with the appropriate combination of methods (feature selection, dimension reduction, and clustering), could provide sufficient information to capture a portion of variance related to the cognitive domain. The importance of FLAIR imaging in predicting MoCA scores is consistent with the previous literature correlating cerebral integrity and potential frailty with cognitive decline and suboptimal recovery post-stroke^51^, attributed to diminished neural plasticity beyond the lesion site^52^. Recently, a research group linked microstructural and macrostructural biomarkers from the normal-appearing brain matter in FLAIR images (texture, intensity, volume) to cognitive function^52^, supporting this hypothesis. In exploring the most predictive motor domain, it is noteworthy that motor outcome predictions uniquely benefit from incorporating additional data derived from tracts and atlases. This observation aligns well with foundational research, highlighting the critical role of structures now known as the corticospinal tract in motor impairment^53^.

In contrast to post-stroke motor impairments that are visible and have been studied exhaustively over the last decades, higher-order symptoms such as depression or anxiety have only recently been recognized and included in the clinical assessment for stroke patients. Thus, prediction approaches used for motor impairment studies cannot simply be translated to other functions yielding the same results. In our analysis, HAD-Depression, and HAD-Anxiety marked the most difficult outcomes to predict with the maximum evaluation scores of R^2^_HAD-D_ =0.034 and R^2^_HAD-A_=0.138. Being able to similarly predict emotional outcomes is crucial, given that one in three stroke survivors is affected by post-stroke depression^54^. Unfortunately, cognitive alterations and mood disorders are frequently overlooked, while they are associated with suboptimal recovery, increased risk of a further stroke, decreased quality of life, and mortality^55,56^. The importance of depression as an independent predictor for functional long-term stroke outcome was already established over two decades ago^57^. Hence, psychological outcomes following a stroke represent a crucial component for further exploration and prediction that are likely to benefit from more complex clinic-radiological models that include additional variables (e.g., genetic vulnerability, biological markers, psychosocial and economic upbringing) to capture and explain more variance.

Additionally, more than one recipe could predict cognitive and psychological scores with comparable accuracy (see Supplementary Figure. 4, Supplementary Table 5). These distinct patterns of feature combinations can be attributed to different factors associated with the same impairment. This demonstrates that the ’optimal recipes’, defined as the most efficient or effective solution under a given set of conditions we have identified are not static but amenable to refinement with integrating a broader array of inputs and methods.

With the novel analytical framework selected for this study, we were able to shed light on the observed and estimate the non-observed feature combinations, suggesting avenues for future investigations. Our score predictions exceeded the predictive accuracy of several comparable (long-term and out-of-sample predicting) attempts reported in the literature^6,53,58^. However, obtaining direct comparisons with existing models following similar objectives is challenging due to the oftentimes not fulfilled criteria of ‘long-term-prediction’ and ‘out-of-sample testing’. While some studies report impressive predictive accuracies for motor scores and functional recovery following stroke, the results reflect a short-term 3-month prediction^59–62^. Supporting the difficulty of comparison, the majority of stroke outcome prediction studies do not employ validation on a previously withheld testing set^63–67^, which is indispensable to claim generalizability and demonstrate the robustness of the models. Our approach focuses on an out-of-sample evaluation to avoid overestimation of the performance (i.e. overfitting), which often occurs in within-sample evaluation or cross-validation methods and do not reflect the true predictive capability of unseen data. In cases where the criteria are fulfilled, the models aim to predict severe complications and mortality after acute ischemic stroke instead of detailed functional, cognitive, or emotional outcomes^68,69^. The assessment and prognosis in these domains typically require comprehensive, time-consuming evaluations by multidisciplinary teams, including neurologists, neuropsychologists, and rehabilitation specialists – leading to limited availability of such data.

However, we also acknowledge that our insights are constrained by the scope of the applied methodologies and clinical data’s utilization (and availability). Here, our primary attempt was to demonstrate the contributing power of neuroimaging modalities for accurate long-term stroke outcome predictions. Therefore, it remains crucial to advocate for embracing open-science principles in the community, and we have made the training dataset available to encourage others to evaluate their predictive models against our cohort. By starting a crowd-sourcing initiative, we are committed to improving stroke outcome predictions and encouraging participation from other teams (to participate or test your algorithm, download the dataset on our website [http://neuralcup.bcblab.com]). The interplay of features helpful in predicting stroke outcomes of diverse domains warrants deeper investigation, and collaboration, methodological exchanges, and data-sharing in the field will greatly advance our understanding.

While investigating predictive inputs and methods is a landmark in stroke outcome research, this study acknowledges its constraints. First and foremost, the quality of the prediction is dependent on the specificity of the behavioral and cognitive assessment. In the present study, while the behavioral assessment might be considered standard neurological practice, the granularity of the cognitive measures might have hampered the predictions. Future sharing initiatives should provide a larger dataset with a more in-depth cognitive examination. Embedding the teams’ observed approaches into the UMAP framework is an innovative step in visualizing potential feature combinations that were directly observed. However, the selection of features included is inherently restricted by the provided data. While we had close to 300 patients with homogeneous data that included acute imaging as well as a chronic behavioral follow-up, we must acknowledge that comparisons to the high number of combinations of features limit the sample size^70^. Additionally, the results reflect the methodologies of 24 prediction submissions, disproportionately influenced by the highest-performing team, thus not representing the full spectrum of possible outcomes. In order to fully assess model performance and predict outcomes in new cohorts, these results should ideally be replicated in the future in a larger population derived from multiple centres, ensuring the generalization of the predictions and allowing us to deploy optimised predictive software as a community. It is essential to provide a wider range of input modalities, such as education level, and employ distinct methodological frameworks to capture a more comprehensive picture. For this, we have created an initiative that facilitates wider participation, increasing the sample size and subsequently aiming to refine the method categories into more nuanced classifications. This could unravel a finer detail of advantageous and suboptimal feature combinations and methodologies for stroke outcome prediction. Another constraint could be raised regarding the selection of neuropsychological test scores and the depth of reported results. Although the UMAP analysis was performed on the complete dataset comprising 13 score predictions (when including the subscores of MOCA), detailed results were only elaborated for five (FM, MoCA, IST, HAD-A, and HAD-D). This was motivated by the comprehensive interpretability of these selected scores across motor, cognitive, and psychological (emotional) impairments and their clinical relevance backed by validated thresholds aiding in predictive model verification and comparison across the field.

Although the study presents certain limitations in quantifying the methodological approaches, it is a pioneering systematic effort to incorporate complete methodologies, exhausting all available variables. This research lays the groundwork for future investigations to enhance the accuracy of stroke outcome predictions. Having demonstrated the viability of out-of-sample predictions, we actively encourage contributions to refine our collective understanding and predictive accuracy for stroke outcomes.

This consortium has established a new benchmark in out-of-sample stroke outcome prediction. We successfully compared the long-term (one-year post-stroke) forecasts across three distinct domains, surpassing prior documented results. Our findings reveal that a universal prediction strategy for stroke outcomes is less effective than employing tailored approaches for each domain or score to achieve the most accurate predictions, holding the promise to improve healthcare for stroke survivors. Our newly proposed analytical framework offers a systematic way to compare different predictive models, laying the groundwork for future benchmarking initiatives and ultimately improving healthcare for stroke survivors.

## Funding

This work is supported by HORIZON- INFRA-2022 SERV (Grant No. 101147319), “EBRAINS 2.0: A Research Infrastructure to Advance Neuroscience and Brain Health”, by the European Union’s Horizon 2020 research and innovation programme under the European Research Council (ERC) Consolidator grant agreement No. 818521 (DISCONNECTOME), the University of Bordeaux’s IdEx ‘Investments for the Future’ program RRI ‘IMPACT’, and the IHU ‘Precision & Global Vascular Brain Health Institute – VBHI’ funded by the France 2030 initiative (ANR-23- IAHU-0001), and OHBM (Organisation for Human Brain Mapping). CR and AT were supported by the NIH of the USA (RF1-MH133701 and P50-DC01466).

## Competing interests

The authors declare that the research was conducted in the absence of any commercial or financial relationships that could be construed as a potential conflict of interest.

## Notes

https://storage.googleapis.com/bcblabweb/neuralcup.html

### Competing Interest Statement

The authors have declared no competing interest.

### Summary of Updates

Added clarification on data-subset selection, modified discussion to ensure the clarity of the aim of the study.

