## Supplementary_Information for "A global effort to benchmark predictive models and reveal mechanistic diversity in long-term stroke outcomes"

**Supplementary Material**

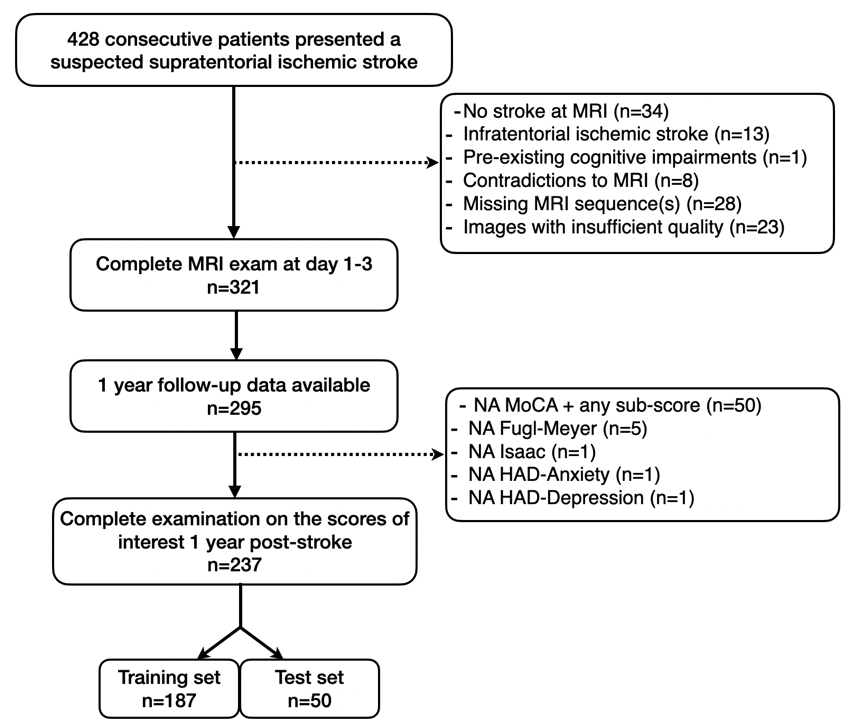

**Supplementary Figure 1.** Flowchart for the sub-dataset used in the NEURAL2023. NA: non-available.

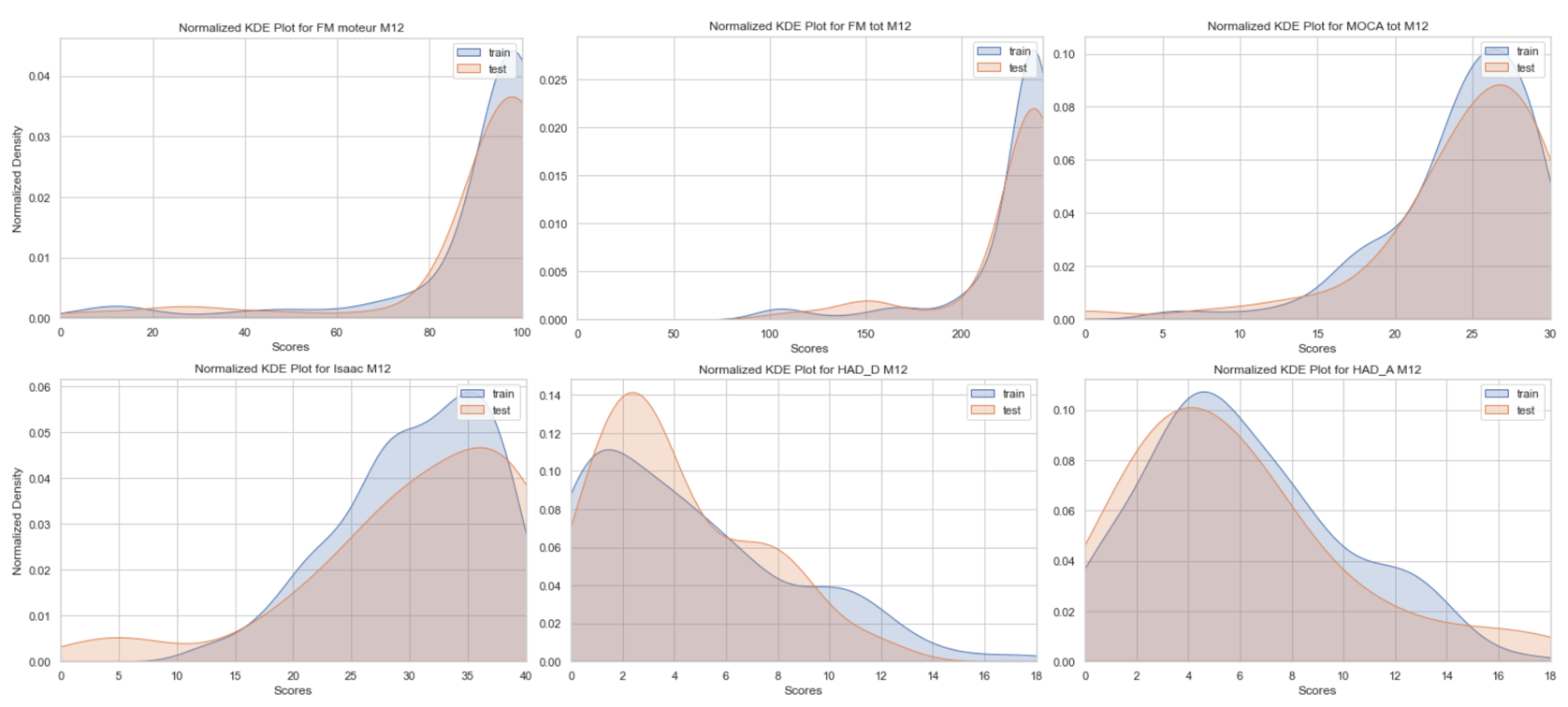

**Supplementary Figure 2.** Normalized Kernel Density Estimation (KDE) Plots for the scores FM_motor, FM_total, MoCA, IST, Had-A, and HAD-D visualizing the largely overlapping distributions of the continuous training and test set variables.

**Supplementary Figure 3.** ROC Curves for the five best predicting teams on a specific score.

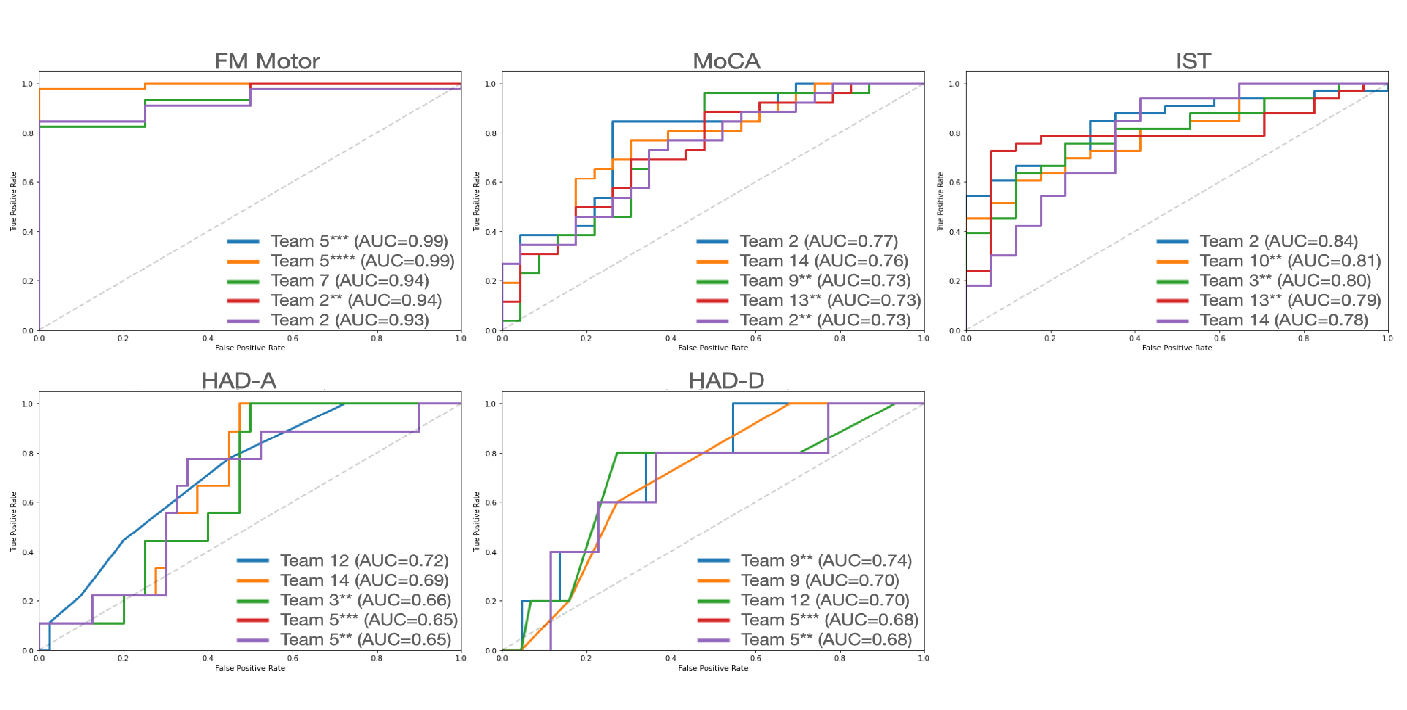

ROC Curves for the five best predicting teams

**Supplementary Table 1.** Methodological Approaches.

|  |  | | **Method** | | **Description** | | **Example*** |
| --- | --- | --- | --- | --- | --- | --- | --- |
|  | **Producing predictions** | | **Artificial Neural Networks (ANNs)** | | Computational models consisting of interconnected nodes organized in layers, which are applied together to process and learn from data; showing the presence of key components such as nodes, layers (input layer, hidden layers, output layer), weights and biases, and an activation function | | neural network model ‘multi-layer perceptron regressor’; convolutional neural networks with multiple hidden layers |
|  |  |  | **Regression Models** | | Analysis of the relationship between a dependent variable and one or more independent variable(s) | | Support Vector Machines; Gradient Boosting Trees Regression; unregularized regression; elastic regression, ridge regression |
|  | **Data extraction/ representation** | | **Feature Selection** | | additional ways to select a subset of relevant features for the prediction model (if any additional analyses involving evaluation of model performance were implemented); if feature selection was embedded into the model (e.g., L2 regularization in ridge regression), the feature was not considered as used by the team | | selection of useful features based on MSE; stability selection using partial least squares variables importance |
|  |  |  | **Clustering** | | Techniques used to group together data into clusters, so that the datapoints within the same cluster are more similar to each other than to those in other clusters | | hierarchical clustering; k-means |
|  |  |  | **Dimension Reduction** | | Reducing the data complexity (e.g., number of input variables in a dataset), while retaining as much information as possible | | Aggregation of T1 signal intensity quantiles; PCA; translating a 3D image into 2D |
|  |  |  | **Parcellation/Segmentation** | | Based on atlases, division of the brain into distinct regions or parcels for the purpose of analyses (e.g., only taking into account a specific brain region to use as model input) | | AAL3 segmentation (anatomical parcellation atlas); white matter resting state atlas; white matter atlas; gray matter atlas; Lesion Quantification Toolkit atlas |
| **Prediction validation/ generalization** | | **Cross Validation** | | Evaluation of the performance and the generalizability of the model by partitioning the training data into subsets, from which some are used for training and some for validation | | nested cross validation, k-folds cross validation | |
|  |  | **Bootstrapping** | | Statistical resampling technique to estimate the distribution of a sample statistic by repeatedly sampling with replacement from the original data | | Stability selection with 250 bootstraps to compute feature reliability and apply thresholds | |

*****From the approaches described by the participating teams.

**Supplementary Formula 1.** Mean Absolute Error.

$$MAE= \frac{1}{n}\sum_{i=1}^{n} |y_{i}-\hat{y}_{i}|$$

*Note:* Formula for MAE, where *n* is the number of observations in the dataset, $y_{i}$ is the true value, and $\hat{y}_{i}$ is the predicted value.

**Supplementary Formula 2.** Mean Squared Error.

$$MSE= \frac{1}{n}\sum_{i=1}^{n} {(y_{i}-\hat{y}_{i})}^{2}$$

*Note:* Formula for MSE, where *n* is the number of observations in the dataset, $y_{i}$ is the true value, and $\hat{y}_{i}$ is the predicted value.

**Supplementary Table 2:** Detailed results of all participating teams’ predictions based on R². The number of the prediction is indicated in the parentheses.

| **Team/Test** | **MOCA** | **IST** | **FM total** | **FM motor** | **HAD-A** | **HAD-D** |
| --- | --- | --- | --- | --- | --- | --- |
| 1 | 0,01351 | 0,02316 | 0,13986 | 0,15688 | 0,0442 | 0,01069 |
| 2 | 0,12275 | 0,34134 | 0,13644 | 0,00056 | 0,08066 | 0,00074 |
| 2 (second) | 0,1356 | 0,33093 | 0,22993 | 0,38935 | 0,0017 | 0,00469 |
| 3 | 0,05581 | 0,03288 | 0,21143 | 0,2771 | 0,02127 | 0,01484 |
| 3 (second) | 0,12786 | 0,16645 | 0,27794 | 0,27211 | 0,13763 | 0,03134 |
| 4 | 2,96E-05 | 0,00183 | 0,0793 | 0,02263 | 0,04104 | 0,00644 |
| 5 | 0,11564 | 0,07834 | 0,54122 | 0,61031 | 0,06247 | 0,00074 |
| 5 (second) | 0,21302 | 0,26255 | 0,14462 | 0,18761 | 0,0573 | 0,01506 |
| 5 (third) | 0,30151 | 0,35293 | 0,54122 | 0,61031 | 0,0573 | 0,00523 |
| 5 (fourth) | 0,30151 | 0,26255 | 0,14462 | 0,18761 | 0,0573 | 0,00523 |
| 6 | 0,0964 | 0,19652 | 0,06736 | 0 | 0 | 0,00897 |
| 7 | 0,03343 | 0,00372 | 0,26613 | 0,44306 | 0,00192 | 0,00045 |
| 8 | 0,01788 | 0,00829 | 0,06574 | 0,08828 | 0,00672 | 7,50E-05 |
| 9 | 0,07174 | 0,07969 | 0,41296 | 0,26276 | 0,00088 | 0,03353 |
| 9 (second) | 0,04781 | 0,02058 | 0,01836 | 0,04348 | 0,00025 | 0,01379 |
| 9 (third) | 0,16928 | 0,12189 | 0,13951 | 0,05954 | 0,02671 | 0,01291 |
| 10 | 0,00229 | 0,01222 | 0,0155 | 0,00205 | 0,00747 | 0,02976 |
| 10 (second) | 0,13121 | 0,27921 | 0,26407 | 0,22439 | 0,1039 | 5,27E-08 |
| 11 | 0,07917 | 0,00191 | 0,17619 | 0,29507 | 0,02978 | 0,01542 |
| 12 | 0,10334 | 0,14672 | 0,04777 | 0,02022 | 0,01815 | 0,0081 |
| 13 | 0,00648 | 0,10759 | 0,11906 | 0,2121 | 3,5E-05 | 0,02496 |
| 13 (second) | 0,14485 | 0,2963 | 0,23523 | 0,22099 | 0,05248 | 0,02844 |
| 14 | 0,16802 | 0,13746 | 0,29525 | 9,76E-05 | 0,0853 | 0,00038 |
| 15 | 0,13496 | 0,06262 | 0,18748 | 0,21928 | 0 | 0 |

**Supplementary Table 3**: Detailed results of all participating teams’ predictions on MoCA sub-scores based on R². The number of the prediction is indicated in the parentheses.

| **Team/Test** | **Visuospatial** | **Denomination** | **Attention** | **Language** | **Abstraction** | **Rappel** | **Orientation** |
| --- | --- | --- | --- | --- | --- | --- | --- |
| 1 | 0,00047 | 0,06724 | 0,04578 | 0,05266 | 0,01446 | 0,00506 | 0,07708 |
| 2 | 0,03519 | 0,0137 | 0,22405 | 0,11657 | 0,07207 | 0,03065 | 0,00575 |
| 2 (second) | 0,00545 | 0,53908 | 0,17441 | 0,0292 | 0,24319 | 0,12556 | 0,00913 |
| 3 | 0,02326 | 0,19019 | 0,06504 | 0,09289 | 0,16818 | 0,00737 | 0,00361 |
| 3 (second) | 0,01791 | 0,18252 | 0,1043 | 0,07346 | 0,19457 | 0,02837 | 0,01426 |
| 4 | 0,00151 | 0,02171 | 0,13146 | 0,01186 | 0,00033 | 0,06497 | 0,00111 |
| 5 | 0,0152 | 0,06568 | 0,22441 | 0,10874 | 0,03333 | 0,00767 | 0,04066 |
| 5 (second) | 0,02745 | 0 | 0,23311 | 0 | 0,09617 | 0,0119 | 0,0911 |
| 5 (third) | 0,0906 | 0,00179 | 0,23426 | 0,09606 | 0,14625 | 0,00139 | 0,00079 |
| 5 (fourth) | 0,0906 | 0,00179 | 0,23311 | 0,00484 | 0,09617 | 0,0119 | 0,0005 |
| 6 | 0,02559 |  | 0,03447 | 0,07435 | 0,0249 | 0,12895 | 0,08815 |
| 7 | 0,00145 | 0,04116 | 0,18179 | 0,20873 | 0,22386 | 0,07279 | 0,00829 |
| 8 | 0,01203 | 0,00163 | 0,00393 | 0,08973 | 0,05076 | 0,00616 | 0,04319 |
| 9 | 0,03891 | 0,13889 | 0,26267 | 0,09376 | 0,02133 | 0,01523 | 0,02758 |
| 9 (second) | 8,14E-05 | 0,20449 | 0,14663 | 0,02743 | 0,0967 | 0,00521 | 0,00833 |
| 9 (third) | 0,01842 | 0,24252 | 0,41999 | 0,12416 | 0,01058 | 0,01308 | 0,06912 |
| 10 | 0,00229 | 0,00142 | 0,0031 | 0,00164 | 0,1119 | 0,07807 | 0,04832 |
| 10 (second) | 0,06013 | 0,01879 | 0,12984 | 0,14577 | 0,11137 | 0,03089 | 0,02164 |
| 11 | 0,01351 | 0,00055 | 0,113 | 0,00248 | 0,14846 | 0,00847 | 0,01743 |
| 12 | 0,01892 | 0,00019 | 0,00684 | 0,01589 | 0,00726 | 0,00802 | 0,05738 |
| 13 | 0,00646 | 0,07279 | 0,03227 | 0,01343 | 0,10135 | 0,0003 | 0,00036 |
| 13 (second) | 0,03863 | 0,15996 | 0,10506 | 0,05504 | 0,17045 | 0,05397 | 0,04292 |
| 14 | 0,00295 | 0,00703 | 0,12172 | 0,23893 | 0,01018 | 0,05568 | 0,03875 |
| 15 | 0 | 0 | 0 | 0 | 0 | 0 | 0 |

**Supplementary Table 4:** Complete Table with ‘optimal recipes’ for all predicted scores for each domain.

| **FEATURE** | **FM** | **MOCA** | **HAD-A** | **HAD-D** | **IST** |
| --- | --- | --- | --- | --- | --- |
| age | 1 | 0 | 0 | 0 | 1 |
| gender | 1 | 0 | 0 | 0 | 1 |
| DWI | 0 | 0 | 0 | 0 | 0 |
| T1 | 1 | 0 | 0 | 0 | 0 |
| segmented lesions | 1 | 1 | 1 | 1 | 1 |
| FLAIR | 0 | 1 | 0 | 0 | 0 |
| tracts | 1 | 0 | 0 | 0 | 0 |
| parcellation atlases | 1 | 0 | 0 | 0 | 0 |
| disconnectomes | 0 | 0 | 1 | 1 | 0 |
| clustering | 1 | 1 | 0 | 0 | 0 |
| ANNs | 1 | 0 | 1 | 1 | 0 |
| regression | 0 | 0 | 1 | 1 | 1 |
| feature selection | 1 | 1 | 0 | 0 | 0 |
| dimension reduction | 0 | 1 | 0 | 0 | 1 |
| parcellation | 0 | 0 | 0 | 0 | 0 |
| cross validation | 0 | 0 | 0 | 0 | 1 |
| *bootstrapping* | 0 | 0 | 0 | 0 | 0 |

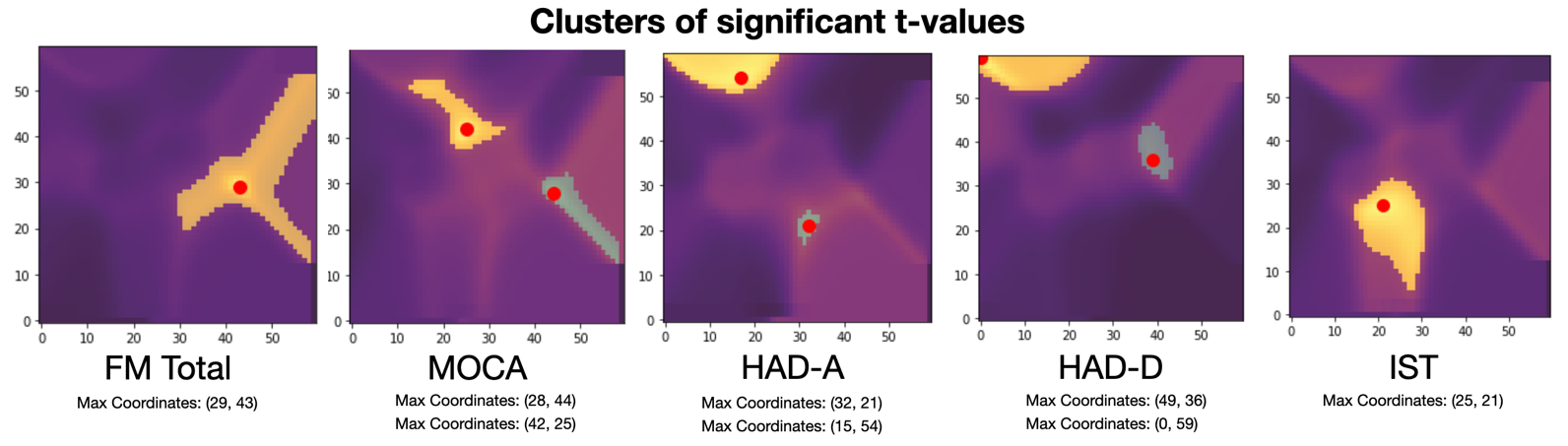

**Supplementary Figure 4.** Clusters of significant t-values. For MoCA, HAD-A, and HAD-D the number of significant clusters is two, indicating that there are two different ‘optimal recipes’ that can potentially work for predicting these scores.

(40, 36)

**Supplementary Table 5:** ‘Optimal Recipes’ for the second significant cluster of the scores MoCA, HAD-A, HAD-D. See the corresponding clusters in Supplementary Figure 4 colored in green.

| **FEATURE** | **MoCA (2)**  coordinates (42, 25) | **HAD-A (2)**  coordinates (32, 21) | **HAD-D (2)**  coordinates (40, 36) |
| --- | --- | --- | --- |
| age | 1 | 1 | 1 |
| gender | 1 | 1 | 1 |
| DWI | 0 | 0 | 0 |
| T1 | 1 | 1 | 0 |
| lesions | 1 | 1 | 1 |
| FLAIR | 0 | 0 | 0 |
| tracts | 0 | 1 | 0 |
| atlases | 0 | 1 | 0 |
| discos | 0 | 0 | 1 |
| clustering | 0 | 1 | 0 |
| ANNs | 1 | 0 | 0 |
| regression | 1 | 0 | 1 |
| feature selection | 1 | 1 | 1 |
| dimension reduction | 0 | 0 | 0 |
| parcellation | 0 | 0 | 1 |
| cross validation | 0 | 0 | 0 |
| *bootstrapping* | 0 | 0 | 0 |

**Supplementary Table 6:** Mean average error (MAE) and mean squared error (MSE) for team and test. The prediction number of each team is represented by the asterisk.

| **Domain** | **Motor** | | **Cognitive** | | | | **Emotional** | | | |
| --- | --- | --- | --- | --- | --- | --- | --- | --- | --- | --- |
| **Test** | **FM Motor** | | **MoCA** | | **IST** | | **HAD-A** | | **HAD-D** | |
| **Team/Error** | **MSE** | **MAE** | **MSE** | **MAE** | **MSE** | **MAE** | **MSE** | **MAE** | **MSE** | **MAE** |
| 1 | 373,23 | 12,76 | 21,91 | 3,75 | 82,65 | 7,30 | 18,22 | 3,29 | 8,90 | 2,48 |
| 2 | 76081,79 | 51,39 | 19,08 | 3,31 | 62,61 | 6,31 | 19,92 | 3,56 | 11,22 | 2,83 |
| 2** | 249,92 | 10,95 | 21,11 | 3,44 | 57,83 | 5,72 | 21,72 | 3,59 | 48,39 | 4,62 |
| 3 | 306,65 | 10,40 | 20,76 | 3,79 | 81,13 | 6,93 | 17,72 | 3,27 | 9,50 | 2,64 |
| 3** | 310,33 | 10,10 | 19,16 | 3,49 | 72,00 | 6,26 | 15,49 | 3,16 | 9,23 | 2,63 |
| 4 | 1651,30 | 33,70 | 90,43 | 7,29 | 132,26 | 9,14 | 26,39 | 4,35 | 29,94 | 4,02 |
| 5 | 226,69 | 8,80 | 22,87 | 3,73 | 79,34 | 6,78 | 16,75 | 3,34 | 10,39 | 2,77 |
| 5** | 329,63 | 10,75 | 18,25 | 3,50 | 68,41 | 6,36 | 16,87 | 3,36 | 9,29 | 2,57 |
| 5*** | 226,69 | 8,80 | 14,72 | 2,99 | 56,70 | 5,99 | 16,87 | 3,36 | 9,65 | 2,43 |
| 5**** | 329,63 | 10,75 | 14,72 | 2,99 | 68,41 | 6,36 | 16,87 | 3,36 | 9,65 | 2,43 |
| 6 | 406,95 | 12,53 | 20,12 | 3,57 | 70,92 | 6,65 | 17,94 | 3,31 | 10,30 | 2,77 |
| 7 | 235,23 | 9,57 | 29,07 | 3,87 | 132,29 | 7,84 | 33,78 | 4,29 | 10,67 | 2,65 |
| 8 | 755,70 | 17,35 | 30,92 | 4,39 | 117,35 | 8,38 | 24,22 | 4,02 | 14,57 | 3,21 |
| 9 | 302,44 | 10,32 | 21,27 | 3,51 | 78,46 | 6,62 | 19,80 | 3,71 | 10,51 | 2,63 |
| 9** | 406,59 | 14,69 | 20,58 | 3,35 | 84,09 | 7,41 | 20,94 | 3,86 | 15,21 | 3,45 |
| 9*** | 493,36 | 11,79 | 17,69 | 3,16 | 74,36 | 6,65 | 17,95 | 3,34 | 9,02 | 2,46 |
| 10 | 491,71 | 12,66 | 22,61 | 3,80 | 93,30 | 7,49 | 20,35 | 3,74 | 11,45 | 3,06 |
| 10** | 317,43 | 11,22 | 19,05 | 3,41 | 65,65 | 6,31 | 18,48 | 3,65 | 13,21 | 3,18 |
| 11 | 331,57 | 13,53 | 23,03 | 3,79 | 95,55 | 7,42 | 18,19 | 3,28 | 17,79 | 3,37 |
| 12 | 739,44 | 18,68 | 54,86 | 5,76 | 160,08 | 10,56 | 21,94 | 3,65 | 11,37 | 2,67 |
| 13 | 333,93 | 10,87 | 21,06 | 3,77 | 75,89 | 6,83 | 18,43 | 3,40 | 11,08 | 2,83 |
| 13** | 327,64 | 10,96 | 19,14 | 3,45 | 64,23 | 6,13 | 17,12 | 3,39 | 9,45 | 2,67 |
| 14 | 457,36 | 12,01 | 22,34 | 3,71 | 72,45 | 6,25 | 17,08 | 3,38 | 15,84 | 3,24 |
| 15 | 326,70 | 11,41 | 18,98 | 3,48 | 78,86 | 6,91 | 17,93 | 3,31 | 9,18 | 2,63 |
